# Effect of behavioral conditions on silk characteristics in the Indian meal moth (*Plodia interpunctella*)

**DOI:** 10.1101/2022.10.11.511611

**Authors:** Toshita V. Barve, R. Keating Godfrey, Caroline G. Storer, Akito Y. Kawahara

## Abstract

Lepidopteran silks are produced during the larval stage and are used for mobility and protection from predators, parasitoids, and pathogens. Our knowledge of silk structure and production in Lepidoptera is based largely on the biology of the domestic silk moth (*Bombyx mori*), but recent comparative evidence suggests that silk production and structure vary widely across moth taxa. Some species like the Indian meal moth (*Plodia interpunctella*) are becoming important biological models to study silk for its potential application to materials science and medicine, but many aspects of silk production in this species remain unknown. Here we characterize the silk of *P. interpunctella* by measuring the width of wandering and pupal silk strands and find that pupal silk is significantly thicker than the latter. We then report individual variation in pupal silk production in our lab-reared colony with a very small number of individuals forgoing pupal silk (< 4%) and find that overcrowding had no effect on this, whereas exposure to elevated temperatures reduced rates of pupal silk production.

## Introduction

Silk is produced by all arthropod subphyla (Sehnal & Sutherland, 2008). Within Hexapoda, where silk gland morphology and silk protein composition vary widely, silk production has arisen and been lost numerous times. Silk production and composition are highly variable across taxa with functions ranging from reproduction and mobility to prey capture and defense. While many hemimetabolous insects produce silk in the adult stage, in holometabolous groups, silk is produced primarily during the larval stage (Sutherland et al., 2010). Lepidoptera larvae produce silk exclusively and use it in many ways, including to fasten themselves to substrate, quickly repel away from danger, and to construct cocoons for protection during metamorphosis (Sourakov and Chadd, 2022).

Insect silk is produced by specialized ectodermal cells and stored in the labial glands, dermal glands, and Malpighian tubules. Lepidoptera larvae create and store silk in the posterior and middle section of the labial gland as an aqueous solution or gel of proteins (Sehnal and Sutherland, 2008). Each strand of silk is composed of two filaments that are made up of a heavy-chain (> 350 kDa) and a light-chain (~25 kDa) fibroin and a glycoprotein held together by sericins, the latter of which are serine-rich proteins that act as a glue (Akai et al., 1987; Mondal et al., 2007). Larvae create a thread of silk by placing a droplet of the silk gel on a substrate, and pulling away from it, thereby stretching the silk proteins into a thread that solidifies as hydrogen bonds form among proteins upon contact with air. The specific amino acid sequence of a silk protein governs the hydrogen bonds formed within and between silk strands and determines secondary structure and silk strength (Fedič et al., 2002; Sutherland et al., 2010).

Because silk has been used as a textile for humankind, a significant amount of research has been conducted on a single species, the domestic silk moth (*Bombyx mori,* Linnaeus, Bombycidae (e.g., Chung et al., 2015; Dong et al., 2016; Mondal et al., 2007). However, interests in silks for a variety of materials sciences purposes has increased in recent years. Other lepidopterans, such as the luna moth (*Actias luna,* Linnaeus, Saturniidae; Chen et al., 2012; Reddy and Yang, 2012), and the Indian meal moth (*Plodia interpunctella,* Hübner, Pyralidae; Milutinović et al., 2020; Shim and Lee, 2015) are now being studied as the search for tougher, stronger, or more flexible silks (Yoshioka et al., 2019). While recent studies on *P. interpunctella* have focused on it as a model organism for genomic study of silk production and evolution (Kawahara et al., 2022), still little is known about physiological properties of silk, variables driving phenotypic plasticity, and how behavioral and environmental variables impact silk production (Harrison et al., 2012; Heryanto et al., 2022; Roberts et al., 2020; Tang et al., 2017).

The Indian meal moth is not only an important moth for silk research, but a significant pest of stored grain. Larvae consume grain products and in the process deposit frass and produce extensive silk galleries that are a contamination risk to these commodities (Mohandass et al. 2007). Since insecticides cannot be readily added to products intended for human consumption, non-invasive environmental approaches to control *P. interpunctella* are of great interest. Surprisingly, chilling larvae to 4 °C can reduce or eliminate pupal silk production (Shim and Lee 2015). While short-term exposure to temperatures exceeding 40 °C can kill larvae (Johnson et al., 2003), why some individuals do not produce silk, and how moderate temperature change can impact silk production is remains unknown. Here we tested whether larval silk varies depending on the function of the silk produced and the environmental conditions of larvae. Specifically, we examined how wandering and pupal silk vary in width and how the presence of conspecifics or increased temperature affects the proportion of individuals that forgo silk production.

## METHODS

### Colony care

All experiments in this paper were performed using a lab population of *P. interpunctella* that was started at the USDA-ARS in Gainesville, FL, USA (see Shirk et al., 2022 for further information on the history of this colony). In 2018, several individuals were transferred to the Florida Museum of Natural History, where a new colony was started for this project. These moths were kept in an airtight 1L Tupperware container placed within a 12 x 12 x 24 in flight cage with a 12/12h light/dark cycle and fed a diet of wheat bran. Temperature and humidity were kept constant, at 21.7 °C and 50.1%, respectively. Gravid females were transferred to an oviposition jar for 24 hours, approximately every 5 weeks to obtain eggs. The following day, eggs were collected and transferred to a container with fresh food so that larvae could be reared.

### Silk collection

Wandering larvae were identified in the rearing container and transferred individually into 2 oz plastic cups with a small amount of food (three trials, N = 55). Larvae were checked every 2-3 days to remove frass and add food as needed. Head widths were measured and wandering silk collected from final instar larvae. In a few instances, multiple head width measurements were taken from individuals for verification. Measurements with a headwidth < 0.75 mm were considered penultimate larvae and excluded from analyses. All other measurements were averaged over the individual for analysis. To collect wandering silk, a larva was placed on a glass microscope slide (Fisher Scientific, 12-552-5) and allowed to wander for one to two minutes or until enough silk had accumulated on the slide. Slides were covered with a #1.5 coverslip (Fisher Scientific, 12544G) held in place with a small amount of clear nail polish (Sally Hansen, Hard as Nails®) at the coverslip corners. To take head width measurements, each larva was sedated in a 1.5 mL PCR tube on ice for two minutes and digital calipers (Neiko Tools, 01407A) were used to measure the widest portion of the head capsule under a dissecting microscope. Cups were cleared of all food and frass to ensure pupal silk was clear of food particles. After pupation, pupal silk was collected by cutting the silk away from the puparium and mounting it on a glass slide with a #1.5 coverslip as described above. Of the 55 larvae, wandering silk was collected from 42 larvae and pupal silk was collected from 16, with both wandering and pupal silk collected from 15 individuals. Data from 6 penultimate instars were excluded from analyses.

### Silk measurement

Silk was imaged using a Leica DM6B microscope system equipped with a Leica DMC6200 camera (Leica Microsystems Inc., Buffalo Grove, IL). Each slide was imaged once, but we used a “stacking” approach to ensure at least 5 strands of silk were in focus (multifocus images were stacked on the z-axis). In cases where stacking could not be accomplished, a single image was taken with as many strands in focus. All images were taken at 20x magnification except for one image that was taken at 10x magnification. Silk strand widths were measured using ImageJ (Schindelin et al., 2012). First, using the region of interest (ROI) manager, a transect line was drawn from the image’s upper left corner to the bottom right corner. Beginning with the top of the image, the first five strands of silk that intersected this line were measured for its width using the line segment tool in ImageJ. For images where fewer than 5 strands were encountered on the first transect, a second transect was drawn from near the first (upper right corner) to the bottom left corner and additional strands were measured until 5 different strands could be measured. Each image was measured by two different human observers to reduce bias, and measurements were averaged for analysis.

### Impact of larval density on pupal silk production

To examine the effect of larval density on the rate of pupal silk production, larvae were housed in one of two conditions: individual containment or community containment. Larvae in the “individual” containment treatment were housed individually in 2 oz plastic cups with food. Community containment consisted of 120 larvae that were split into 4 groups of 30 larvae each. Each group was placed into a jar with layered pieces of corrugated cardboard to serve as “houses” for larval pupation. Over the course of two weeks, the presence and absence of pupal silk was documented.

### Effect of elevated temperature on silk production

To test whether elevated temperature affects the proportion of caterpillars producing pupal silk, we exposed caterpillars to one of three treatments: reared at room temperature (RT, 23 - 25 °C, 40 - 50% relative humidity), reared chronically at high temperature in an incubator (IN, 33 - 34 °C, 40 - 50% RH), or transferred to high temperature for acute exposure during the final instar (TR). We established four biological replicates for each condition, with the exception of IN where we established three biological replicates. Unfortunately, the first and second IN replicates did not produce larvae. Thus, we used larvae from the third IN replicate to make technical replicates.

Replicates were set up by sedating adults in the colony with CO_2_ for <1 minute. Eight to ten sedated adults were placed into a small, 8 oz glass jar with food. We repeated this four times so that each replicate represented a different generation from source colonies. Jars were placed in their respective environmental conditions; RT and TR adults were maintained at room temperature and IN adults were provided 24 hours at RT for mating and egg laying before being placed in the incubator. All replicates were checked every other day and food and water were added as needed. Twenty-five wandering larvae were isolated from each replicate, leading to 100 larvae isolated for each environmental condition and a total of 300 isolated larvae. We recorded the presence or absence of pupal silk in each isolation cup once pupation was complete.

### Hypothesis testing and statistics

All data were analyzed in R version 4.1.2 (R Core Team, 2021; RStudio Team, 2022). Observers did not differ significantly in their quantification silk types (wandering silk: chi-squared = 3.67, df = 2, p = 0.1598; pupal silk: chi-squared = 0.458, df = 2, p = 0.7953). We therefore averaged silk width measurements across observers. We test our main hypothesis in two ways: first, we compared all measurements of wandering silk with the smaller sample size of pupal silk measurements using a Mann-Whitney U Test (Wilcoxon Rank Sum Test). Second, we compared widths only from individuals for which we had both wandering and pupal silk measurements. On this subset of samples, we conducted a paired t-test with the null hypothesis being that there is no difference between wandering and pupal silk widths. For experiments testing the role of aggregation and temperature on pupal silk production, pairwise Chi-Square tests of independence were used to detect statistically significant differences in the proportion of caterpillars producing pupal silk in each treatment. The Benjamini-Hochberg procedure with false discovery rate set (FDR) to 0.05 was used to correct for multiple comparisons.

## RESULTS

### Variation in silk width

Silk width did not correlate with head width in last instar larvae (t = - 1.1873, *df* = 34, p = 0.243), suggesting larger larvae do not necessarily produce larger silk strands. Our analysis comparing silk widths between wandering silk and pupal silk revealed a statistically significant difference in the silk produced for these different purposes (Figure 3). Pupal silk is on average over one micrometer thicker than wandering silk (= 3.28, *SD* = 0.64 μm and = 2.12, *SD* = 0.32 μm, respectively; p < 0.0001). Because we collected more width measurements from wandering silk than pupal silk, we compared silk widths within individuals (N = 15). Supporting our initial results, we found that pupal silk width is greater than that of wandering silk produced by the same individual prior to pupating (t = −7.612, *df* = 14, p < 0.0001).

### Effect of environmental conditions on pupal silk production

Pupal silk production varied slightly, but most individuals produced pupal silk (= 96.5, *s* = 4%, 4 replicates, N = 150). Those that did not produce pupal silk and initiated metamorphosis in a “naked” puparium all survived to adulthood, indicating silk is not required for development under lab conditions. Increased larval density did not affect pupal silk production, as individually housed larvae produced pupal silk at a frequency indistinguishable from those housed in groups (p = 0.376, Figure 2A). Temperature, on the other hand, had a noticeable effect on pupal silk production and survival. Larvae transferred to a higher temperature just prior to pupation (TR) and those reared at a higher temperature (IN) both showed significantly lower amount of pupal silk from those reared under ambient conditions (RT = 99%; TR = 93%; IN = 15%; Figure 2B). Additionally, IN treatment larvae suffered significantly greater mortality, and often failed to reach pupation compared to the other two treatments (RT = 1%; TR = 9%; IN = 69%; p < 0.0001 for both pairwise contrasts; Figure 2B). Final instar larvae in the IN treatment reached a diapause-like state, as wandering larvae at room temperature pupated an average of 5 (*s* = 2) days after being isolated, whereas 16 of the 100 isolated larvae in the IN treatment had not pupated for more than 25 days after isolation (Figure 2B).

**Figure 1.**
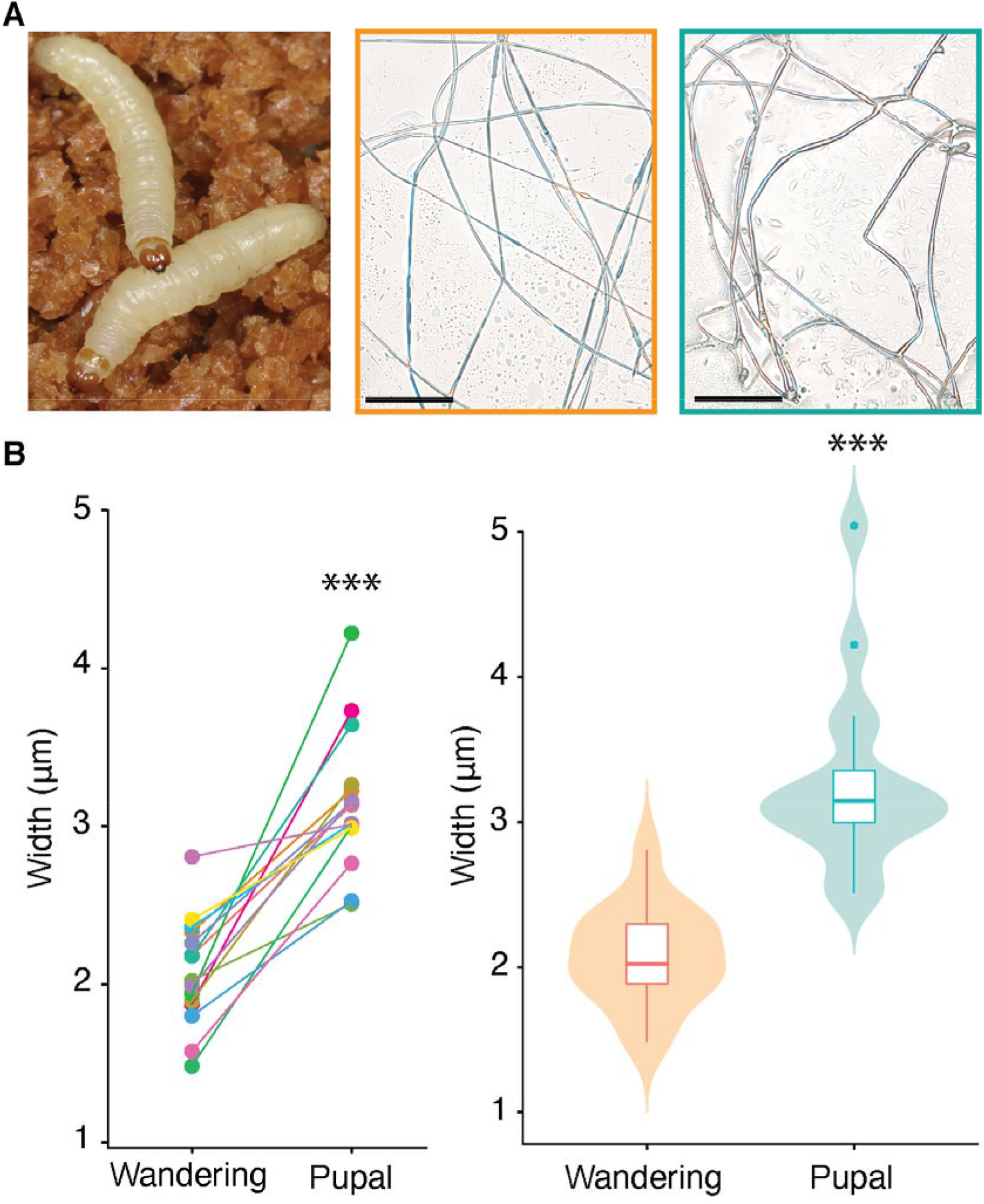
*Plodia interpunctella* and differences in its silk across behavioral conditions. (A) From left to right: wandering larvae with food substrate, wandering silk, pupal silk. Scale bar = 100 μm. (B) Differences in pupal silk width from intra-individual (left) and sample-level (right) comparisons. *** = p < 0.001.

**Figure 2.**
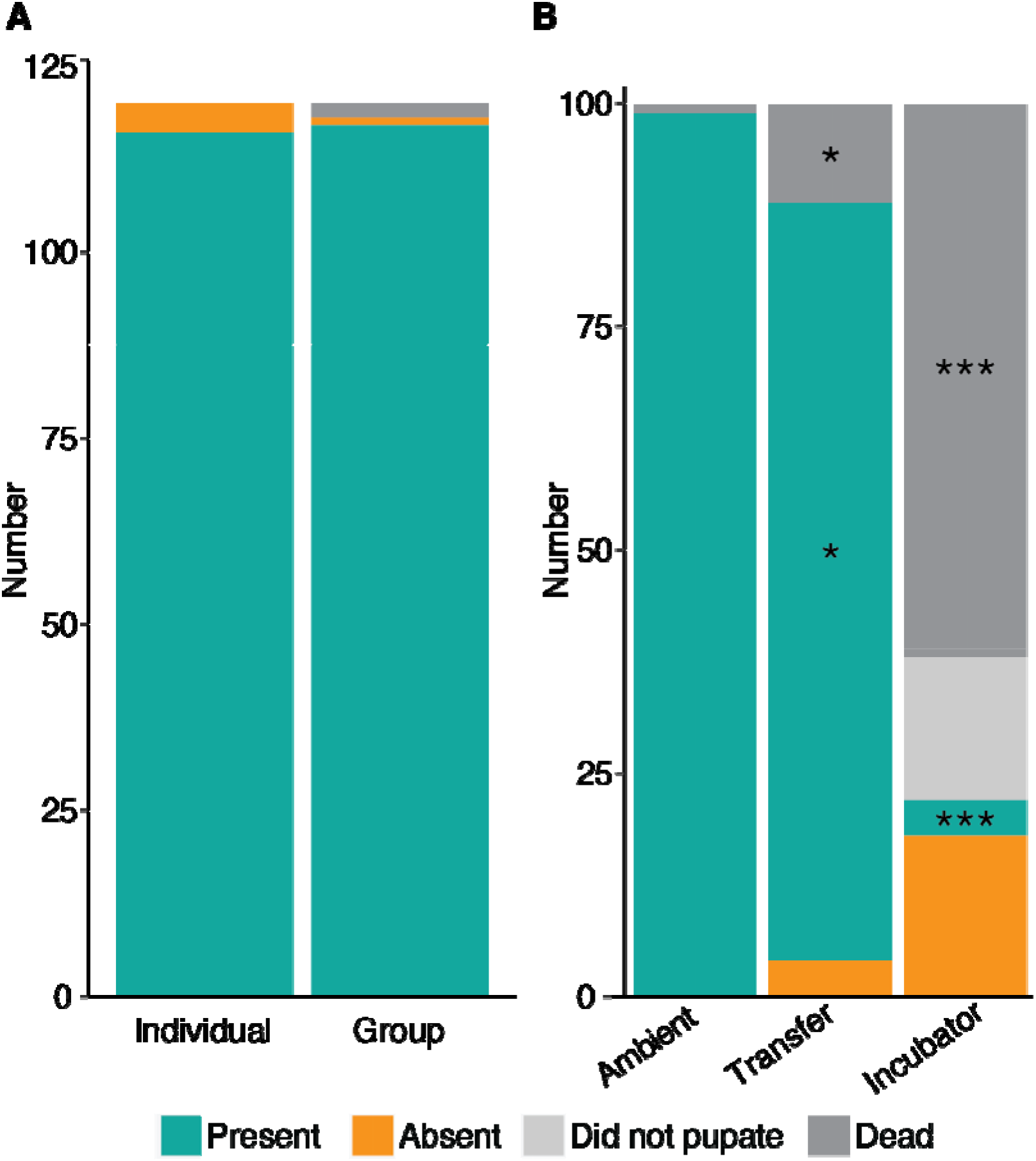
Factors affecting pupal silk production in *P. interpunctella.* (A) No significant difference detected in the proportion of larvae producing pupal silk housed individually or with conspecifics. (B) Higher mortality and lower pupal silk production when larvae are treated with increased temperature, either shortly before pupation (Transfer) or for the duration of larval development (Incubator). Asterisks indicate comparisons with ambient conditions of the same color. * = p < 0.05; *** = p < 0.001.

## DISCUSSION

*Plodia interpunctella* is an important, emerging model organism for silk research, yet factors that impact silk production and structure are not well-characterized. Here we find that wandering and pupal silk vary in width, suggesting that larvae can modify physical characteristics of silk strands based on the silk’s primary function. Our data also show that there can be variation in pupal silks and that higher ambient temperature can reduce pupal silk production.

Silk produced in larval labial glands is a synapomorphy of Amphiesmenoptera, the superorder that includes Lepidoptera and Trichoptera (Sehnal and Sutherland, 2008; Yonemura et al., 2009), a clade that dates back to more than 200 mya (Misof et al. 2014). Indeed, the heavy- and light-chain fiborins that comprise the primary silk proteins produced by Amphiesmenoptera are highly conserved (Yonemura et al., 2009). There is also an extensive overlap in the ecological use of silk among Lepidoptera and caddisflies, with larvae using silk for mobility and as well as protection from predators and parasitoids in both orders. Pupal silk in particular is important for protecting Lepidoptera during metamorphosis, as it provides a means of securing the puparium in cavities or crevices, may provide camouflage, and can act as a physical barrier against parasitoids (Fedič et al., 2002; Lindstedt et al., 2019). In addition to sericins and fibroins, silk contains antimicrobial seroin proteins that are added during the construction of silk fibers and act to prevent infections (Singh et al., 2014). Predation and parasitism at the pupal stage can be quite high for moths, with estimates of mortality often above 60% for species that show cyclic population densities (East 1974; Stark & Harper 1982; Lindstedt, Murphy, & Mappes 2019). Lindstedt et al. (2019) hypothesized that pupal silk may be thicker than wandering silk, as pupae cannot escape and need a protective barrier against predators. We found that individual pupal silk strands are wider in *P. interpunctella* and may contribute to a thicker layer of silk forming the cocoon. This structural difference could derive from changes in larval spinning behavior or differences in protein composition. In *B. mori,* seroin proteins are accumulated in labial glands of the final instar and released only once the pupal silk creation commences (Dong, Song, et al. 2016), suggesting pupal silk protein composition may be different than wandering silk.

Interestingly, at least two parasitoid species are attracted to silk compounds (Ha et al., 2006), suggesting that there may be tradeoffs to silk production. While we found no evidence for the presence of conspecifics affecting variation in pupal silk production, temperature significantly reduced the proportion of larva that made pupal silk. Chronic exposure to elevated temperatures near 33 °C throughout larval development resulted in severe mortality and diapause. While low temperatures are known to induce diapause (Johnson et al., 1995), to our knowledge elevated temperatures have not been reported to trigger diapause. The shift in developmental timing and elevated mortality made it difficult to confidently compare silk production in the chronically elevated temperature treatment (IN) with acute exposure condition (TR) or controls (RT). Larvae that experienced an acute temperature increase after most of their larval development was complete did not suffer as severe mortality, but still showed a significant reduction in pupal silk production, suggesting that environmental factors can affect this behavior. However, while temperature affects many aspects of larval biology, including development, physiology, and immune function (Catalán et al., 2012; Fontenot et al., 2012; Na & Ryoo, 2000), temperature can also affect physical properties of silk. Shim & Lee (2015) report that chilling larvae can result in solidification of silk glands, likely rendering them nonfunctional. Therefore, it is possible that rather than inducing a physiological or behavioral change in larvae, higher temperatures may affect physical properties of silk after silk strands have been synthesized.

## Acknowledgements

We are grateful to Paul Shirk for providing the moths that were used to start our colony in 2018. We thank Rhys Campo, Christian Couch, Hailey Dansby, and Amanda Markee for their assistance with colony maintenance and silk width data collection. Paul Frandsen and David Plotkin helped improve the quality of the manuscript. This project was supported in part by the Florida Museum of Natural History, the University Scholars Program run by the Center for Undergraduate Research at the University of Florida, University of Florida Research Opportunity Fund (DRPD-ROF2019) and NSF EF-2217159.

## Literature Cited

Akai, H., Imai, T., & K. Tsubouchi. 1987. Fine-structural changes of liquid silk in the silk gland during the spinning stage of Bombyx larvae. J. Sericultural Sci. Jpn. 56, 131–137. https://doi.org/10.11416/kontyushigen1930.56.131

Catalán, T.P., Wozniak, A., Niemeyer, H.M., Kalergis, A.M., & F. Bozinovic. 2012. Interplay between thermal and immune ecology: Effect of environmental temperature on insect immune response and energetic costs after an immune challenge. J. Insect Physiol. 58, 310–317. https://doi.org/10.1016/j.jinsphys.2011.10.001

Chen, F., Porter, D., & F. Vollrath. 2012. Structure and physical properties of silkworm cocoons. J. R. Soc. Interface 9, 2299–2308. https://doi.org/10.1098/rsif.2011.0887

Chung, D.E., Kim, H.H., Kim, M.K., Lee, K.H., Park, Y.H., & I.C. Um. 2015. Effects of different *Bombyx mori* silkworm varieties on the structural characteristics and properties of silk. Int. J. Biol. Macromol. 79, 943–951. https://doi.org/10.1016/j.ijbiomac.2015.06.012

Dong, Z., Zhao, P., Zhang, Y., Song, Q., Zhang, X., Guo, P., Wang, D., & Q. Xia. 2016. Analysis of proteome dynamics inside the silk gland lumen of *Bombyx mori.* Sci. Rep. 6, 1–10. https://doi.org/10.1038/srep21158

Fedič, R., Žurovec, M., & F. Sehnal. 2002. The silk of Lepidoptera. J. Insect Biotechnol. Sericology 71 71, 1–15. https://doi.org/10.11416/jibs2001.71.1

Fontenot, E.A., Arthur, F.H., Nechols, J.R., & J.E. Throne. 2012. Using a population growth model to simulate response of *Plodia interpunctella* Hübner to temperature and diet. J. Pest Sci. 85, 163–167. https://doi.org/10.1007/s10340-011-0411-0

Ha, D.M., Choi, S.H., Shim, J.K., Jung, D.O., Song, K.S., Nho, S.K., & K.Y. Lee. 2006. Behavioral attraction of two parasitoids, *Venturia canescens* and *Bracon hebetor*, to silk extracts of a host *Plodia interpunctella.* J. Asia-Pac. Entomol. 9, 287–292. https://doi.org/10.1016/S1226-8615(08)60305-2

Harrison, P.W., Mank, J.E., & N. Wedell. 2012. Incomplete sex chromosome dosage compensation in the Indian meal moth, *Plodia interpunctella*, based on *de novo* transcriptome assembly. Genome Biol. Evol. 4, 1118–1126. https://doi.org/10.1093/gbe/evs086

Heryanto, C., Hanly, J.J., Mazo-Vargas, A., Tendolkar, A., & A. Martin. 2022. Mapping and CRISPR homology-directed repair of a recessive white eye mutation in *Plodia* moths. iScience 25, 103885. https://doi.org/10.1016/j.isci.2022.103885

Johnson, J.A., Wang, S., & J. Tang. 2003. Thermal death kinetics of fifth-instar *Plodia interpunctella* (Lepidoptera: Pyralidae). J. Econ. Entomol. 96, 519–524. https://doi.org/10.1093/jee/96.2.519

Johnson, J.A., Wofford, P.L., & R.F. Gill. 1995. Developmental thresholds and degree-day accumulations of Indianmeal moth (Lepidoptera: Pyralidae) on dried fruits and nuts. J. Econ. Entomol. 88, 734–741. https://doi.org/10.1093/jee/88.3.734

Kawahara, A.Y., Storer, C.G., Markee, A., Heckenhauer, J., Powell, A., Plotkin, D., Hotaling, S., Cleland, T.P., Dikow, R.B., Dikow, T., Kuranishi, R.B., Messcher, R., Pauls, S.U., Stewart, R.J., Tojo, K., & P.B. Frandsen. 2022. Long-read HiFi sequencing correctly assembles repetitive heavy fibroin silk genes in new moth and caddisfly genomes. Gigabyte 2022, 1–14. https://doi.org/10.46471/gigabyte.64

Lindstedt, C., Murphy, L., & J. Mappes. 2019. Antipredator strategies of pupae: How to avoid predation in an immobile life stage? Philos. Trans. R. Soc. B Biol. Sci. 374. https://doi.org/10.1098/rstb.2019.0069

Milutinović, M., Čurović, D., Nikodijević, D., Vukajlović, F., Predojević, D., Marković, S., & S. Pešić. 2020. The silk of *Plodia interpunctella* as a potential biomaterial and its cytotoxic effect on cancer cells. Anim. Biotechnol. 31, 195–202. https://doi.org/10.1080/10495398.2019.1575848

Misof, B., Liu, S., Meusemann, K., Peters, R.S., Donath, A., Mayer, C., Frandsen, P.B., Ware, J., Flouri, T., Beutel, R.G., Niehuis, O., Petersen, M., Izquierdo-Carrasco, F., Wappler, T., Rust, J., Aberer, A.J., Aspöck, U., Aspöck, H., Bartel, D., Blanke, A., … & X. Zhou. 2014. Phylogenomics resolves the timing and pattern of insect evolution. Science 346(6210), 763–767. https://doi.org/10.1126/science.1257570

Mohandass, S., Arthur, F.H., Zhu, K.Y., & J.E. Throne. 2007. Biology and management of Plodia interpunctella (Lepidoptera: Pyralidae) in stored products. J. Stored Prod. Res. 43 (3): 302–311.

Mondal, M., Trivedy, K., & S.N. Kumar. 2007. The silk proteins, sericin and fibroin in silkworm, *Bombyx mori* Linn., A review. Casp. J. Environ. Sci. 16.

Na, J.H., & M.I. Ryoo. 2000. The influence of temperature on development of *Plodia interpunctella* (Lepidoptera: Pyralidae) on dried vegetable commodities. J. Stored Prod. Res. 36, 125–129. https://doi.org/10.1016/S0022-474X(99)00039-9

Reddy, N., &. Y. Yang. 2012. Investigation of the structure and properties of silk fibers produced by *Actias lunas.* J. Polym. Environ. 20, 659–664. https://doi.org/10.1007/s10924-012-0482-x

Roberts, K.E., Meaden, S., Sharpe, S., Kay, S., Doyle, T., Wilson, D., Bartlett, L.J., Paterson, S., & M. Boots. 2020. Resource quality determines the evolution of resistance and its genetic basis. Mol. Ecol. 29, 4128–4142. https://doi.org/10.1111/mec.15621

Schindelin, J., Arganda-Carreras, I., Frise, E., Kaynig, V., Longair, M., Pietzsch, T., Preibisch, S., Rueden, C., Saalfeld, S., Schmid, B., Tinevez, J.-Y., White, D.J., Hartenstein, V., Eliceiri, K., Tomancak, P., & A. Cardona. 2012. Fiji: an open-source platform for biological-image analysis. Nat. Methods 9, 676–682. https://doi.org/10.1038/nmeth.2019

Sehnal, F., & T. Sutherland. 2008. Silks produced by insect labial glands. Prion 2, 145–153. https://doi.org/10.4161/pri.2.4.7489

Shim, J.K., Lee, K.Y., 2015. Chilling results in failure of silk secretion by wandering larvae of *Plodia interpunctella*. J. Asia-Pac. Entomol. 18, 483–487. https://doi.org/10.1016/j.aspen.2015.06.003

Shirk, B.D., Shirk, P., Furlong, R.B., Scully, E.D., Wu, K., & B.D. Siegfried. 2022. Gene editing of the ABC Transporter/White locus using Crispr/Cas9-Mediated mutagenesis in the Indian meal moth. SSRN Electron. J. https://doi.org/10.2139/ssrn.4192442

Singh, C.P., Vaishna, R.L., Kakkar, A., Arunkumar, K.P., & J. Nagaraju. 2014. Characterization of antiviral and antibacterial activity of *Bombyx mori* seroin proteins: Accessory silk proteins as antimicrobial agents. Cell. Microbiol. 16, 1354–1365. https://doi.org/10.1111/cmi.12294

Sourakov, A., & R.W. Chadd. 2022. The lives of moths: a natural history of our planet’s moth life. Princeton University Press, Princeton. 288 pp.

Sutherland, T.D., Young, J.H., Weisman, S., Hayashi, C.Y., & D.J. Merritt. 2010. Insect silk: One name, many materials. Annu. Rev. Entomol. 55, 171–188. https://doi.org/10.1146/annurev-ento-112408-085401

Tang, P.-A., Wu, H.-J., Xue, H., Ju, X.-R., Song, W., Zhang, Q.-L., & M.-L. Yuan. 2017. Characterization of transcriptome in the Indian meal moth *Plodia interpunctella* (Lepidoptera: Pyralidae) and gene expression analysis during developmental stages. Gene 622, 29–41. https://doi.org/10.1016/j.gene.2017.04.018

Yonemura, N., Mita, K., Tamura, T., & F. Sehnal. 2009. Conservation of silk genes in Trichoptera and Lepidoptera. J. Mol. Evol. 68, 641–653. https://doi.org/10.1007/s00239-009-9234-5

Yoshioka, T., Tsubota, T., Tashiro, K., Jouraku, A., & T. Kameda. 2019. A study of the extraordinarily strong and tough silk produced by bagworms. Nat. Commun. 10, 1469. https://doi.org/10.1038/s41467-019-09350-3

